# A data-driven method to identify frequency boundaries in multichannel electrophysiology data

**DOI:** 10.1101/2020.07.09.195784

**Authors:** Michael X Cohen

## Abstract

**Background:** Electrophysiological recordings of the brain often exhibit neural oscillations, defined as narrowband bumps that deviate from the background power spectrum. These narrowband dynamics are grouped into frequency ranges, and the study of how activities in these ranges are related to cognition and disease is a major part of the neuroscience corpus. Frequency ranges are nearly always defined according to integer boundaries, such as 4-8 Hz for the theta band and 8-12 Hz for the alpha band.

**New method:** A data-driven multivariate method is presented to identify empirical frequency boundaries based on clustering of spatiotemporal similarities across a range of frequencies. The method, termed gedBounds, identifies patterns in covariance matrices that maximally separate narrowband from broadband activity, and then identifies clusters in the correlation matrix of those spatial patterns over all frequencies, using the dbscan algorithm. Those clusters are empirically derived frequency bands, from which boundaries can be extracted.

**Results:** gedBounds recovers ground truth results in simulated data with high accuracy. The method was tested on EEG resting-state data from Parkinson’s patients and control, and several features of the frequency components differed between patients and controls.

**Comparison with existing methods:** The proposed method offers higher precision in defining subject-specific frequency boundaries compared to the current standard approach.

**Conclusions:** gedBounds can increase the precision and feature extraction of spectral dynamics in electrophysiology data.

## Introduction

Population neural activity, as measured through LFP, EEG, and MEG, exhibits multiple temporal features that have been linked to cognition and disease, including phasic evoked responses (Luck and Kappenman, 2013), broadband fluctuations (Miller et al., 2014, 2009), and narrowband rhythmic activity (Buzsáki and Watson, 2012; Engel et al., 2001; Uhlhaas and Singer, 2006). Narrowband activity has garnered considerable interest over the past few decades, because it provides empirical and theoretical links to behavior, neurophysiology, animal studies, and computational models (Cohen, 2011; Klimesch, 1999; Wang, 2006).

Narrowband activity can be defined as a spectral peak that deviates from the “background” frequency spectrum, which typically exhibits a 1/f-like shape such that spectral power decreases with higher frequencies. Multiple peaks can be observed in EEG spectra and have been given labels such as delta, theta, alpha, beta, and gamma.

Grouping and studying frequencies in ranges is not arbitrary, because neurons and circuits have biophysical properties (e.g., ion channel time-constants) that constrain their timing, thus leading to neural rhythms with particular temporal characteristics (Buzsáki and Wang, 2012; Jones et al., 2009; Wang, 2006). Thus, there are compelling reasons why the neuroscience community adopts the frequency-band-naming conventions.

However, the exact numerical boundaries between frequency ranges are somewhat subjective. For example, the alpha band is variously defined as 7-12 Hz, 8-13 Hz, 9-11 Hz, etc (Niedermeyer, 1997). Others have distinguished subbands such as low vs. high alpha (8-10 Hz vs. 10-12 Hz) (Klimesch, 1999). Sometimes the frequency boundaries are selected a priori and other times they are based on visual inspection of the data. On the one hand, these arbitrary (and nearly always integer-based) cut-offs probably do fairly little damage to the final conclusions of each individual publication. But on the other hand, they add uncertainty and a sense of subjectivity to the literature as a whole. Furthermore, fixed boundaries prevent more nuanced and potentially informative analyses into how these boundaries may differ as a function of group (e.g., patient group or genetic background), or other individual factors such as age, personality, task performance, etc.

Here I propose an empirical method to identify spectral boundaries based on spatiotemporal patterns. The method, termed “gedBounds” for generalized eigendecomposition boundaries, relies on the assumption that all frequencies within the same band have highly similar spatiotemporal characteristics, and thus that boundaries between bands can be defined as sudden changes in those similarities. gedBounds works on individual data, thus allowing both for individually optimized frequencies and for group comparisons (e.g., whether a patient group has statistically significantly different frequency boundaries compared to controls). The method is fast (a few minutes per dataset, depending on the amount of data and desired frequency resolution), deterministic (it produces the same results each time it is run on the same data), and easy to implement (MATLAB code is provided).

## Materials and methods

### Data types and overview of gedBounds

gedBounds is appropriate for multichannel electrophysiology in which narrowband activity can be expected. This includes EEG, MEG, and LFP. The method is designed for analyzing, not for cleaning, and thus cleaned data should be used, with as many artifacts filtered out as possible.

The gedBounds method can be broken down into three stages. These are briefly described below, and detailed over the rest of this section.

1. Create a channel covariance matrix using broadband (non-temporally filtered) data. This matrix is called **R** for reference (Figure 1a).
2. Loop over a range of frequencies (e.g., 2-100 Hz). At each frequency, create a channel covariance matrix using narrowband filtered data. This matrix is called **S** for signal. Compute the generalized eigendecomposition on **S** and **R**, which identifies a spatial filter (a set of weights for all channels) that maximally separates **S** from **R**, while ignoring any features that are common between the two matrices. The eigenvector of maximal separation is stored (Figure 1a).
3. Pairwise squared correlations across eigenvectors from all frequencies are computed and stored in a matrix of eigenvector similarities (Figure 1b). A cluster analysis is applied to this matrix to identify “blocks” on the diagonal that have high similarities (Figure 1b). These clusters are frequency ranges, and their edges are the frequency boundaries (Figure 1c).

### Underlying assumptions

There are two key assumptions underlying the validity of this method. The first and most important assumption is that frequencies of the same band have strongly correlated spatiotemporal dynamics. To give an example, the topographies and time courses of neural activity filtered at 11.0 Hz and at 11.1 Hz are likely to be extremely highly correlated, partly because neural activity is non-stationary and partly because narrowband filters entail some spectral smoothing. However, the topographies and time courses of activity filtered at 11 Hz and at 5 Hz can be expected to correlate weakly or modestly, because these two frequencies fall into different bands and thus have distinct spatiotemporal dynamics. Although in principle the spatiotemporal patterns could transition smoothly from theta to alpha, empirically this is not the case, and there will be some frequency that shows a sharp change in correlation with its neighboring frequency; this will be considered the boundary that separates theta from alpha.

**Figure 1.**
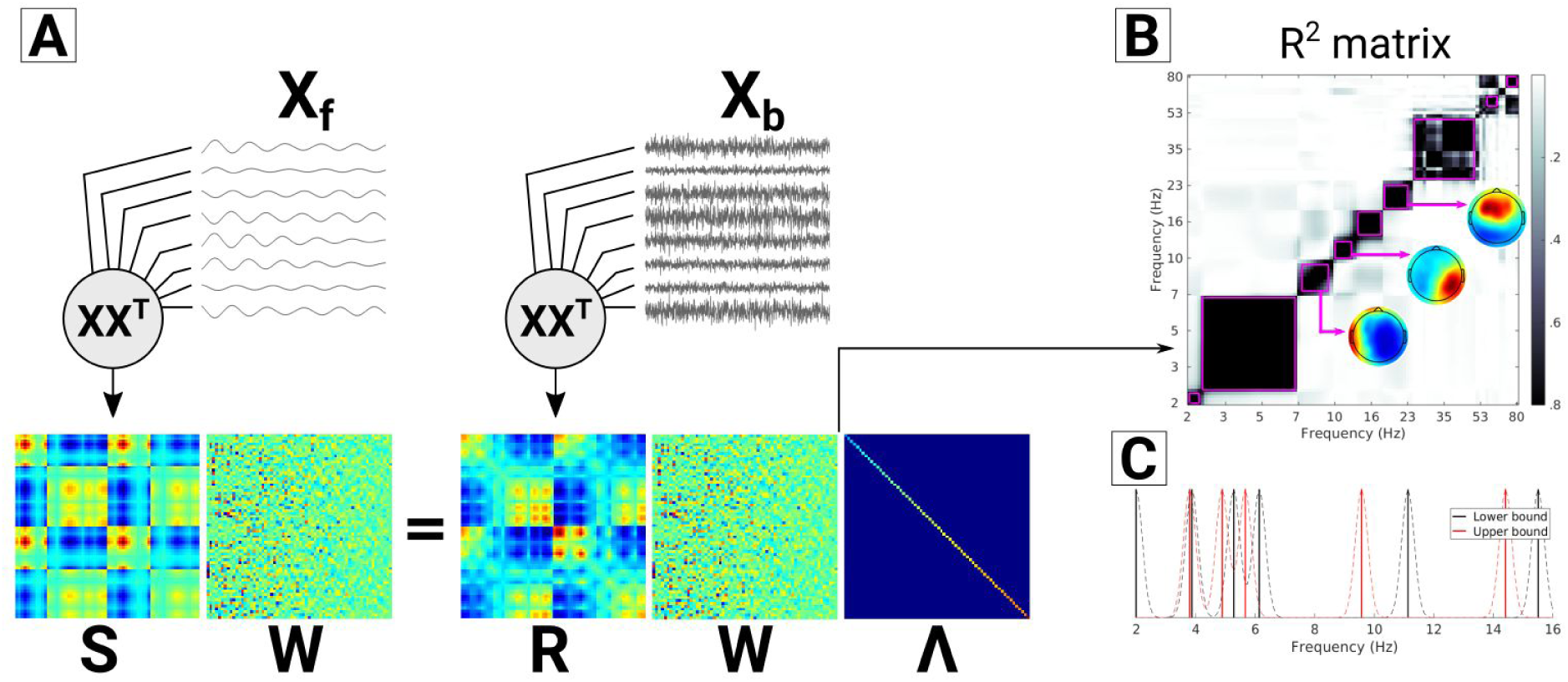
Overview of gedBounds procedure. **A**) Compute channel covariance matrices (**S** and **R**) from broadband signal (**X**_**b**_) and narrowband-filtered signals (**X**_**f**_). The spatial filter that best separates their corresponding covariances is the eigenvector associated with the largest generalized eigenvalue. **B**) The pairwise correlations across eigenvectors from all frequencies are computed, and clustered to identify the diagonal blocks, which are taken as frequency bands. Spatial maps can be visualized from each block. **C**) The boundaries can be visualized on a spectral plot and smoothed with a Gaussian to facilitate cross-subject comparisons.

The second assumption underlying this method is that spatiotemporal characteristics within a frequency change relatively little over time. This is because the method relies on covariance matrices; non-stationary dynamics are averaged together, and thus only the (roughly) stationary and linear components of the channel interactions are preserved in a covariance matrix. In practice, this is a reasonable assumption because neural generators generate stable topographies. Applying this method to task-related designs, in which cognitive events may change rapidly, is discussed in the Discussion section.

### Generalized eigendecomposition

Eigendecomposition is a matrix decomposition that identifies important features, or patterns, in a square matrix, e.g., a covariance matrix. It is the basis for many applications in statistics, including principal components analysis. Generalized eigendecomposition (GED) identifies features that maximally separate two matrices, while ignoring features that are common to both matrices. Correlating eigenvectors across frequencies, rather than correlating the topographical distributions of the power values, has several advantages: GED (1) separates narrowband from broadband activity that are recorded simultaneously, thus holding constant cognitive and other factors; (2) reduces the impact of artifacts or non-brain sources that have a relatively wide frequency distribution; (3) helps suppress noise; (4) takes into account both spatial and temporal dynamics instead of only spatial or only temporal features; (5) has higher signal-to-noise ratio characteristics and is more accurate at recovering ground truth simulations compared to principal or independent components analyses (Cohen, 2017; de Cheveigné and Parra, 2014; Nikulin et al., 2011; Zuure and Cohen, 2020).

The goal of GED is to find a spatial filter (a set of weights used to compute a weighted combination of all electrodes) that maximizes the ratio of the energy in two covariance matrices. Covariance matrices are used because they encode all pairwise linear relationships across the electrodes, thus allowing the method to leverage volume conduction and spatial autocorrelation to identify spatial patterns in the data, as opposed to the reduced signal-to-noise quality of a single electrode. Two covariance matrices are computed: One from the narrowband filtered signal (channels-by-time data matrix **X**_**f**_; obtained from narrowband filtering as described later), and one from the broadband signal (channels-by-time data matrix **X**_**b**_).

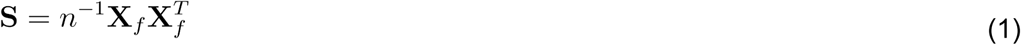

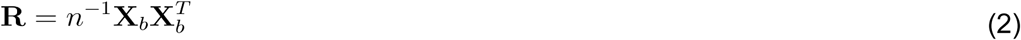

The letters **S** and **R** indicate that these matrices are considered signal and reference. In the empirical data, continuous time series were segmented into 2-second epochs, and the covariance matrix within each epoch was computed. Covariance matrices were cleaned by rejecting epochs with non-representative covariance matrices. The Euclidean distance from each covariance matrix to the epoch-averaged covariance matrix was computed and z-scored, and the average covariance matrix was re-computed using only epochs with covariance matrices within 3 standard deviations away from the mean.

The spatial filter that maximally separates the two covariance matrices is obtained by finding the channel vector **w** that maximizes the Rayleigh quotient:

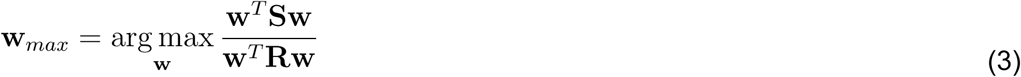

The solution to the finding ***w***_max_ comes from a generalized eigenvalue decomposition on matrices **S** and **R**.

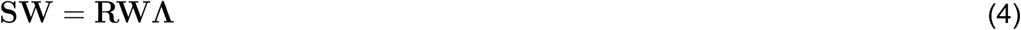

The diagonal elements in matrix **Λ** are the eigenvalues, each of which is the ratio of equation 3 for each corresponding column of **W**. Thus, the column of **W** with the largest associated eigenvalue is the spatial filter that maximizes the multivariate energy ratio between the narrowband and the broadband activity. This procedure is repeated for a range of temporal frequencies (each creating a different **S** matrix) with the same **R** matrix.

A small amount of shrinkage regularization (1%) was applied to the **R** matrix in order to improve the quality of the decomposition. In our experience, 1% shrinkage has no appreciable effect on decompositions of clean, full-rank, and easily separable data, and considerably improves the decompositions of noisy or reduced-rank data. Shrinkage regularization involves shrinking down the covariance matrix and adding to its diagonal a fraction of the average eigenvalues.

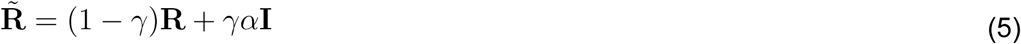

*γ* is the amount of shrinkage (.01) and *α* is the average of all eigenvalues of **R. I** is the identity matrix.

The component time series is obtained by multiplying **w** by the data matrix **X**, although the component time series were not used here. The eigenvectors comprise a mixture of boosting signal and suppressing noise, and are therefore difficult to interpret anatomically. They are the backwards model of “looking through the channels to see the source.” The anatomically interpretable topography (“looking from the source to the channels”) comes from applying the eigenvector to the **S** covariance matrix (Haufe et al., 2014) as **w**^T^**S**. Because there are many eigenvectors for each frequency band, the approach taken here was to perform a principal components analysis (PCA) on the set of eigenvectors or component maps, and report the top PC. This is referred to as PCA-averaging in the text.

### Eigenvectors correlation matrices

Although the component maps are anatomically interpretable, the eigenvectors are dominated by higher spatial frequencies because they partially filter out volume conduction. Thus, the eigenvectors are well suited for distinguishing neighboring narrowband activity patterns because they will be considerably less correlated with each other compared to the component maps.

For this reason, eigenvectors from the top component from each frequency were stored, and then a correlation matrix was computed between all pairs of eigenvectors. Because there is sign-uncertainty in eigenvectors (that is, eigenvectors have no inherent sign as the subspace they identify spans an infinite line crossing the origin), and to facilitate clustering, the correlations were squared.

### Clustering into frequency bins

The aforementioned correlation matrix has a block-diagonal-like pattern. The interpretation is that each block reflects one spatiotemporally coherent frequency band, and thus the goal is to isolate those blocks. This was done using an algorithm called density-based spatial clustering of applications with noise (dbscan) (Ester et al., 1996). Dbscan is a non-hierarchical clustering method that uses thresholding and local concentration to identify clusters in data.

The key parameter of dbscan is the step size for searching in neighborhoods to define each cluster. The parameter is called epsilon and in this application, its units are r^2^ values. Small values of epsilon lead to unacceptably small clusters, whereas large values of epsilon lead to a single cluster spanning multiple frequency bands. The method developed here to identify an appropriate epsilon was inspired by a Bayesian approach of fixing the data and varying the parameter, and then selecting the parameter that maximized the following quality function.

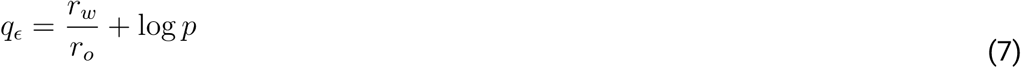

*q*_*ϵ*_ is the quality measure for parameter value *ϵ, r*_*w*_ is the average correlation values inside the clusters and *r*_*o*_ is the average correlations outside the clusters. That ratio can select for very small clusters with very high correlations, and therefore the *log(p)* term was added, where *p* is the proportion (between 0 and 1) of the total correlation matrix occupied by the clusters. That term adds a cost for very small clusters. The two terms together find a balance between increased selectivity and increased cluster size. The epsilon associated with the largest peak in the *q* function was taken as the final value used for the clustering. If there was no clear peak, then the average epsilon across the tested range was selected. The optimal thresholds were close to 0 (average 0.025 in the empirical data), because the correlation matrices tended to be sparse, with most r^2^ values being essentially 0 and a small number being close to 1.

Note that dbscan need not include each frequency in a cluster if the eigenvectors are not sufficiently similar to their neighbors. In other words, it is possible to have, for example, a theta cluster that ends at 8.3 Hz and an alpha cluster that begins at 9.2 Hz.

Many clustering methods exist. Hierarchical clustering, k-means, and t-SNE were also explored here, but dbscan generally gave the most interpretable and sensible results. There are several advantages of dbscan for this application, including that dbscan does not require assigning each frequency to a cluster; “noise” frequencies are identified and ignored; the number of clusters does not need to be defined a priori (unlike k-means, for example); cluster assignment is deterministic; the key parameter (epsilon) is easier to identify compared to the number of clusters; and each frequency is exclusively assigned to only one cluster.

### Identifying frequency boundaries

To compare frequency boundaries across subjects, the boundaries were converted into spectral likelihood functions through a kernel density estimator. For each subject, a vector was constructed with a value of one for the frequency of each boundary, and zeros everywhere else. This impulse vector was convolved with a Gaussian with a 2 Hz full-width at half maximum. These smoothed impulse vectors were then averaged over all subjects.

Frequency bins were quantitatively compared across the two groups by finding the boundaries closest to the standard frequency boundaries (e.g, 8-12 Hz for alpha).

### Temporal filtering

Data were temporally narrowband filtered by convolution with a Morlet wavelet, defined here as a Gaussian in the frequency domain. Extracted frequencies ranged from 2 Hz to 80 Hz in 100 logarithmically spaced steps (logarithmic instead of linear steps were chosen to make the frequency range block sizes comparable across the spectrum). The full-width at half-maximum of the Gaussian varied from 2 to 5 Hz with increasing frequency.

The exact choice of filtering method is not crucial for the successful application of this method; any other suitable narrowband filter could be used. Spanning a wide frequency range from 2-100 Hz is also not necessary; one could restrict the analysis to, e.g., 5-15 Hz to identify possible subbands of alpha (this is illustrated in the Results section). However, it is important to have a high frequency resolution in order to identify precise boundaries. This is particularly important if the goal is to treat the boundaries as dependent variables in a subsequent analysis.

### Simulated data

Simulated data were used to validate the method with ground-truth patterns. The simulation methods were similar to those used elsewhere (Cohen, 2017). Briefly, an EEG forward model was generated using the OpenMEEG package (Gramfort et al., 2010) in the Brainstorm toolbox (Tadel et al., 2011) in MATLAB, comprising 2004 dipoles placed around the cortex and 64 electrodes on the scalp, following the 10-20 system. Projecting time series from each dipole to the scalp and summing over all dipoles provides a biophysically plausible model of EEG signals. Dipole time series were generated as 10 seconds of random broadband Gaussian noise, and the time series from three selected dipoles were generated as narrowband noise (simulating narrowband non-stationary time series). Electrode-level data were then analyzed as described above.

### Empirical data

Data were taken from the PRED+CT database (http://predict.cs.unm.edu/downloads.php) (Cavanagh et al., 2017), accession number d002 (Cavanagh, JF, Kumar, P, Mueller, AA, Richardson, SP, Mueen, A, 2018). Data were recorded from 60 scalp EEG channels during a resting-state period between 1.5 and 10 minutes (average 3.5 minutes). There were 56 subjects in total, half of which were Parkinson’s disease patients and half were matched controls. Only the off-medication session was used here. Note that this dataset was selected based on public availability and convenience; as this is a methods paper, the focus is on the example application rather than possible relevance of the findings to the diagnosis, treatment, or etiology of Parkinson’s disease.

Data preprocessing included high-pass filtering at .5 Hz, attenuating 60 Hz line noise using the ZapLine method (de Cheveigné, 2020), transforming to average reference, and removing noisy data features via independent components analysis with the jade algorithm (Cardoso, 1999). The average number of components removed was 3.1, which was not significantly different between the patient and control groups (t_54_=.83, p=.41). Data were stored in eeglab (Delorme and Makeig, 2004) format. The gedBounds method and code does not rely on the eeglab toolbox, and could easily be adapted to any other data-storage format.

### Code availability

MATLAB code to generate the simulation and implement gedBounds can be found at mikexcohen.com/data/gedBounds.zip. Upon acceptance this zip file will be posted as supplemental material accompanying this publication. The code relies on the Statistics and Machine Learning toolbox for the dbscan() function. Empirical data can be downloaded online as described above.

## Results

### Validation on simulated data

Narrowband signals in three dipoles in the brain were generated and projected to the scalp, along with Gaussian noise at 2001 other dipoles. Figure 2a shows the matrix of r^2^ values from all pairs of eigenvectors. The red dashed lines illustrate the actual frequency boundaries, the solid magenta lines illustrate the empirically derived frequency clusters via dbscan clustering, and the blue line shows the eigenspectrum. Note that with dbscan (unlike, e.g., k-means clustering), the number of clusters is not specified a priori. Thus, the three clusters were automatically identified based on the patterns of correlations, not based on prior knowledge of the true number of frequency ranges.

**Figure 2.**
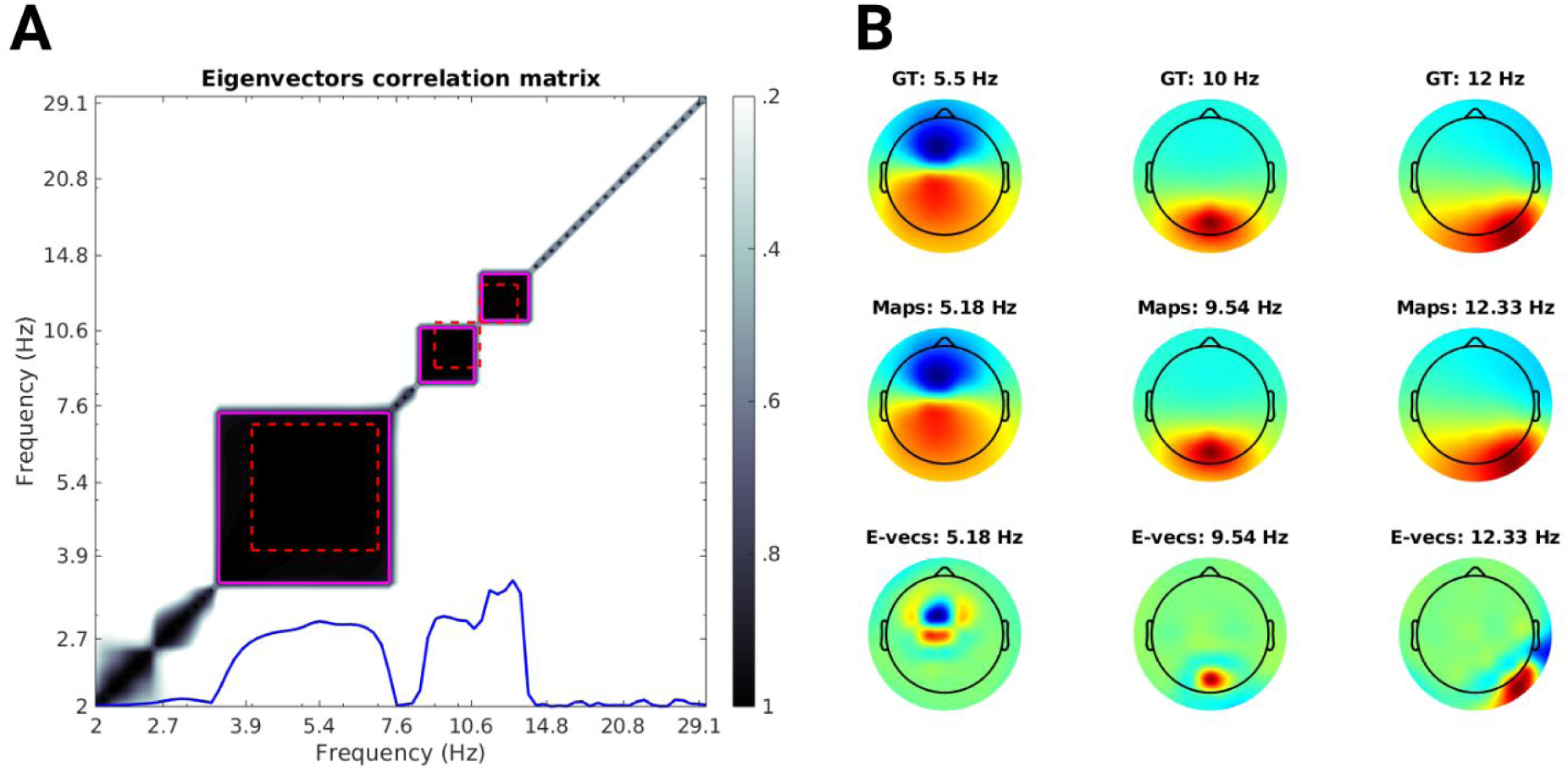
Results from simulated EEG data. **A**) Squared-correlation matrix of all pairs of eigenvectors. The red dashed line shows the true underlying source frequencies, the solid magenta line shows the result of the dbscan clustering, and the blue line on the bottom shows the top eigenvalue per frequency (the measure of separation between the narrowband and broadband covariance matrices). **B**) Topographical maps showing the ground truth dipole projections (“GT”), the component maps PCA-averaged over all frequencies in each cluster, and the PCA-averaged eigenvectors (“E-vecs”) within each cluster.

The empirical clusters are slightly larger than the ground-truth clusters due to spectral leakage from the temporal filtering. Interestingly, however, the boundary between the two “alpha” clusters (9-11 Hz and 11-13 Hz) is sharp, because their eigenvector patterns are uncorrelated. It is interesting to compare this sharp transition to the gentle transition between the two “apha” peaks in the eigenspectrum (blue line in Figure 2a). This shows that clusters in the eigenvectors correlation maps are more sensitive to subtle changes over frequency, compared to the eigenspectrum alone. This is, in fact, not surprising: the eigenvectors encode detailed spatial patterns while the eigenvalues encode scalar separability from the broadband signal.

The topographies of the component maps averaged across all frequencies within these three clusters closely matched the simulated data (Figure 2b). This figure also shows how the component maps can be spatially correlated while the eigenvector correlation matrix shows sharp boundaries. Indeed, the eigenvectors tend to be more spatially localized, as they counter the effects of volume conduction, although they are less anatomically interpretable (Haufe et al., 2014).

When the frequencies overlap across different bands (e.g., 9-11 Hz in one dipole and 10-13 Hz in another dipole), then gedBounds will form clusters around the non-overlapping frequencies. An example is shown in supplementary Figure S1. It is also possible that the components can be better separated by considering later components instead of the largest component (see Discussion section).

Repeating the analysis using PCA on the narrowband covariance matrices instead of GED revealed poor recovery of the ground-truth dynamics (Figure S2). This resulted from the very large r^2^ values across all pairs of frequencies. In this case, PCA is driven by the collective projections of the dipoles, and thus mainly reflects the average of the leadfield model as opposed to frequency-specific activity. The same is likely true for biological data.

Overall, the simulated data show that the gedBounds method is effective and precise at recovering boundaries between neighboring frequencies, under the assumption that the projections of the source onto the electrodes are linearly separable. This performance justifies applying the method to empirical data, in which ground truth is unknown.

### Empirical data: frequency clusters and boundaries

Figure 3 shows results for one dataset. As in the simulated data, strong correlations around the main diagonal of the eigenvectors correlation matrix are expected due to spectral leakage of neighboring frequencies. However, the important feature is the “block-diagonal” appearance, with the correlations suddenly decreasing across neighboring frequencies. This is taken as evidence for spatiotemporally separable frequency bands. Correlation matrices with empirical frequency clusters for all patients and controls are shown in supplementary Figures S3-4.

**Figure 3.**
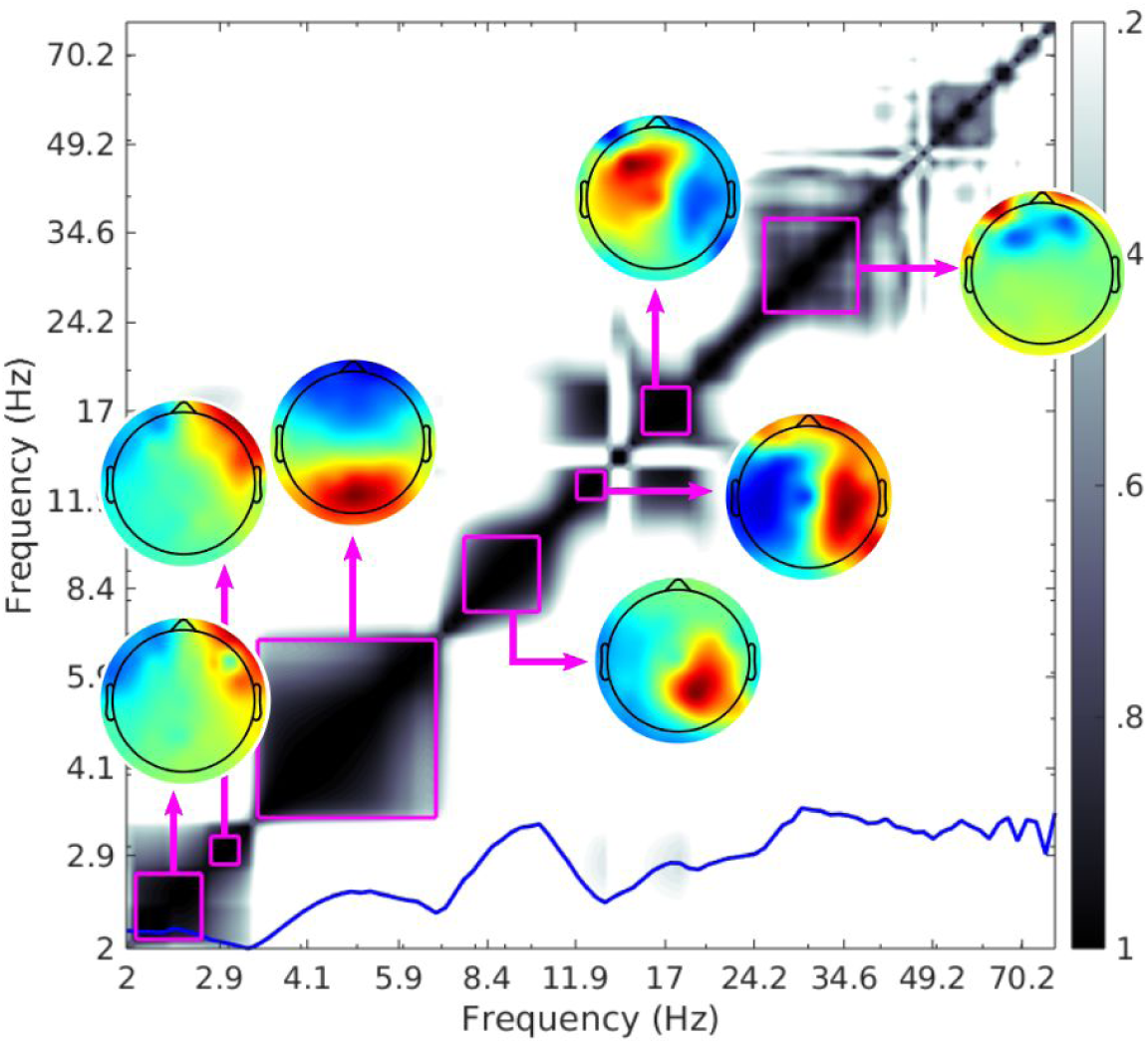
gedBounds results for one individual. Topographical maps show the PCA-averaged forward model for each cluster. The blue line shows the height-normalized eigenvalue spectrum. Note that clusters of high correlation do not always coincide with clear peaks in the eigenvalue spectrum.

The number of frequency bands ranged between 4 and 12 (Figure 4a), and was significantly larger in Parkinson’s patients compared to controls (averages: 8.61 and 6.85, t(54)=3.19, p=.002). Average correlations within a cluster tended to decline with increasing frequency (Figure 4b). In other words, lower frequencies tended to be more spatiotemporally robust within a frequency band. This linear trend was statistically significant in the control group (p=.045) but not in the Parkinson’s group (p=.16). The selection of low correlations at very low frequencies may be due to two separate bands being clustered together, which can be observed in some datasets in the supplemental Figures S3 and S4 (more on this observation below).

**Figure 4.**
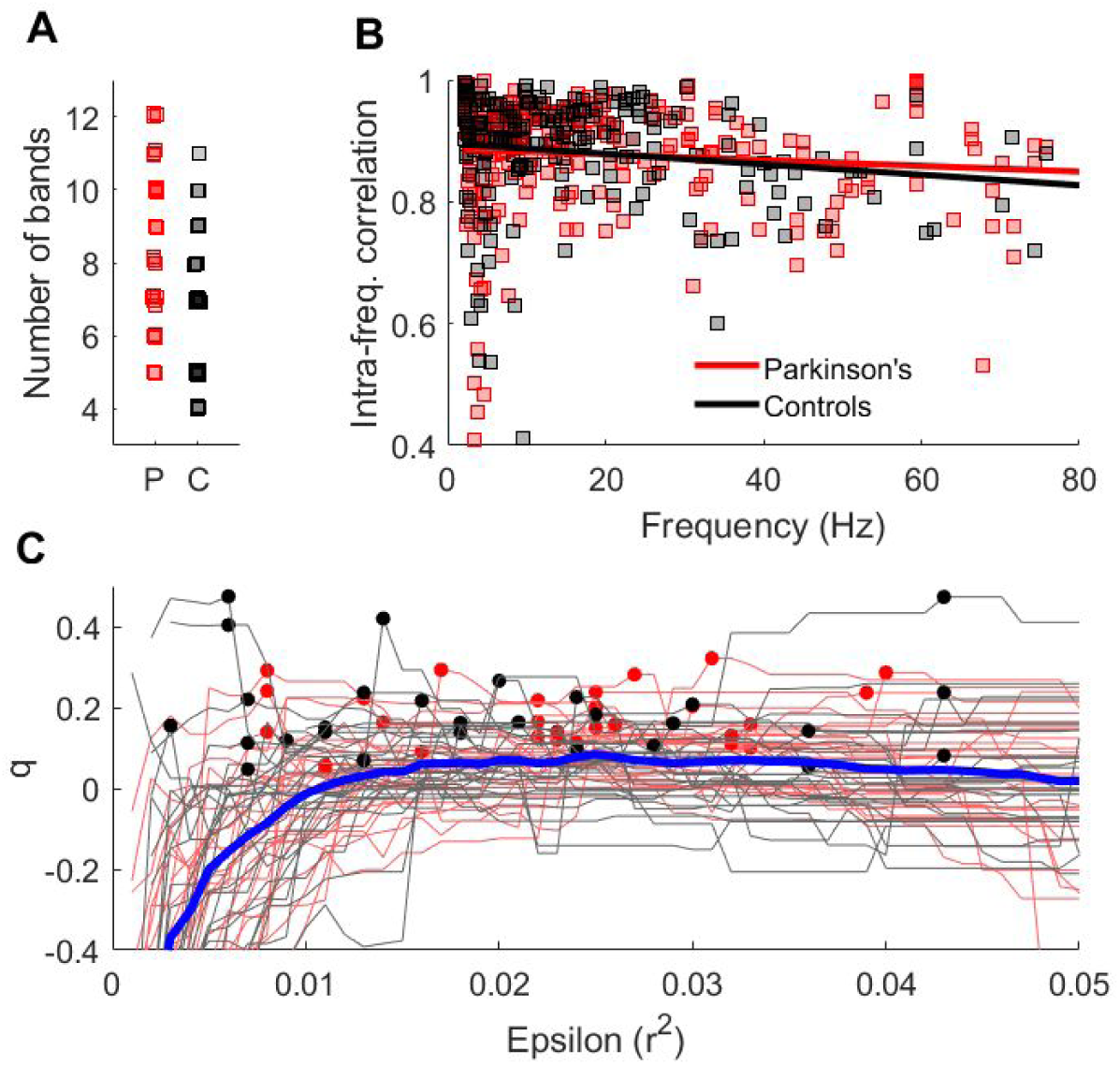
Group-aggregated empirical results. **A**) Number of frequency bands identified by the dbscan clustering (small, random offsets were added for visibility). P = Parkinson’s, C = controls. **B**) The average squared-correlation values within each cluster (essentially the average “blackness” inside each magenta box in Figures S3-4) are plotted as a function of the center frequency of that cluster. Straight lines indicate best-fit linear slopes. **C**) The q vector (equation 7) as a function of the key dbscan parameter epsilon (neighborhood search step size). Each line is an individual and the blue line shows the grand average.

Figure 4c shows the q vector (equation 7) over a range of epsilon values. Each line is from a subject, and the dots indicate the peak parameter selected for that individual. The optimal epsilon parameter was not different between the groups (t(54)=-.619).

Visual inspection of the eigenvector correlation matrices suggests that some clusters could be interpreted as comprising subclusters. Because the number of clusters was defined based on global properties, it could be refined for local investigations. For example, Figure S3 (repeated in Figure 5) seems to suggest that the low frequency band in s811 could be further subdivided. This was done by repeating the same decomposition algorithm with the same parameters, except for frequencies 1-10 Hz instead of 2-100 Hz. Figure 5 shows that using this focused frequency range subdivided the 2-7 Hz cluster into two separate clusters from 1-3.6 Hz and 4-6.4 Hz.

**Figure 5.**
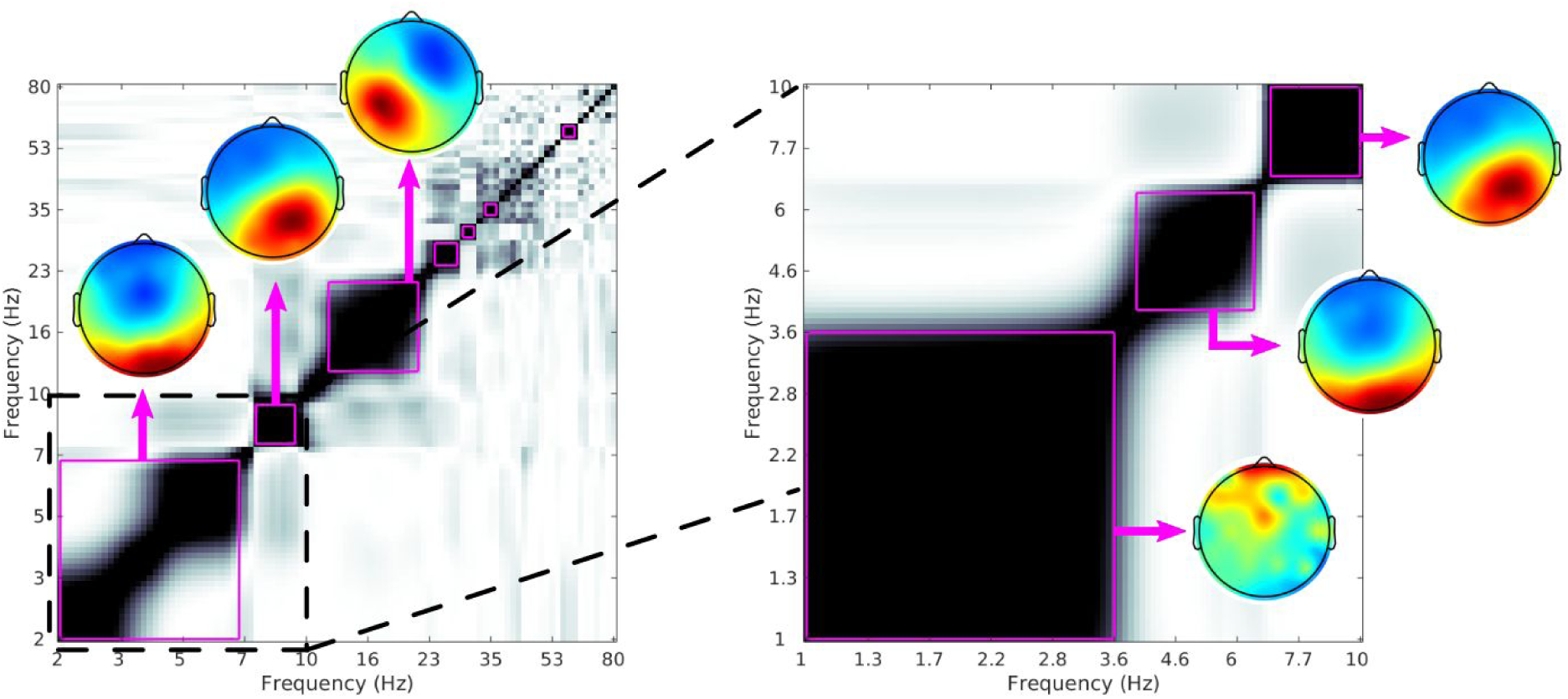
Example of higher-resolution spectral band identification by running gedBounds using a limited frequency range. The left panel shows the algorithm applied to a range of 2-80 Hz (same as Figure S3, box 811). The right panel shows the same algorithm with the same parameters, but applied to data filtered in a range of 1-10 Hz. This increased the sensitivity to detecting lower-frequency subbands. Topographies show the PCA-averaged component maps from all frequencies within each cluster.

### Aggregating boundaries across datasets

To pool data across individuals, a kernel density estimator (KDE) approach was taken, in which the frequency boundaries for each individual were marked as impulses on a frequencies vector, and then that impulse-frequencies vector was smoothed with a 2 Hz FWHM Gaussian (Figure 1c). The motivations for Gaussian-smoothing are (1) it facilitates cross-subject averaging and (2) it reflects some uncertainty in the estimation, in other words, it is interpreted as a likelihood distribution around a true frequency boundary for which the empirical-derived frequency is the maximum likelihood estimate. Finally, the smoothed KDEs are averaged over subjects. Other widths for the Gaussian were explored, from 1 Hz to 5 Hz, and gave qualitatively similar results.

Results are shown in Figure 6. In general, the upper boundaries tended to be more consistent across subjects compared to the lower boundaries. Figure 6c shows the bounds for the alpha band, defined as the lower bound closest to but not exceeding 10 Hz, and its associated upper bound. A two-samples t-test on the average alpha bandwidth (upper minus lower bounds) between the groups was statistically significant (t(53)=2.2, p=.032; one control subject with an alpha bandwidth exceeding 3 standard deviations was excluded).

**Figure 6.**
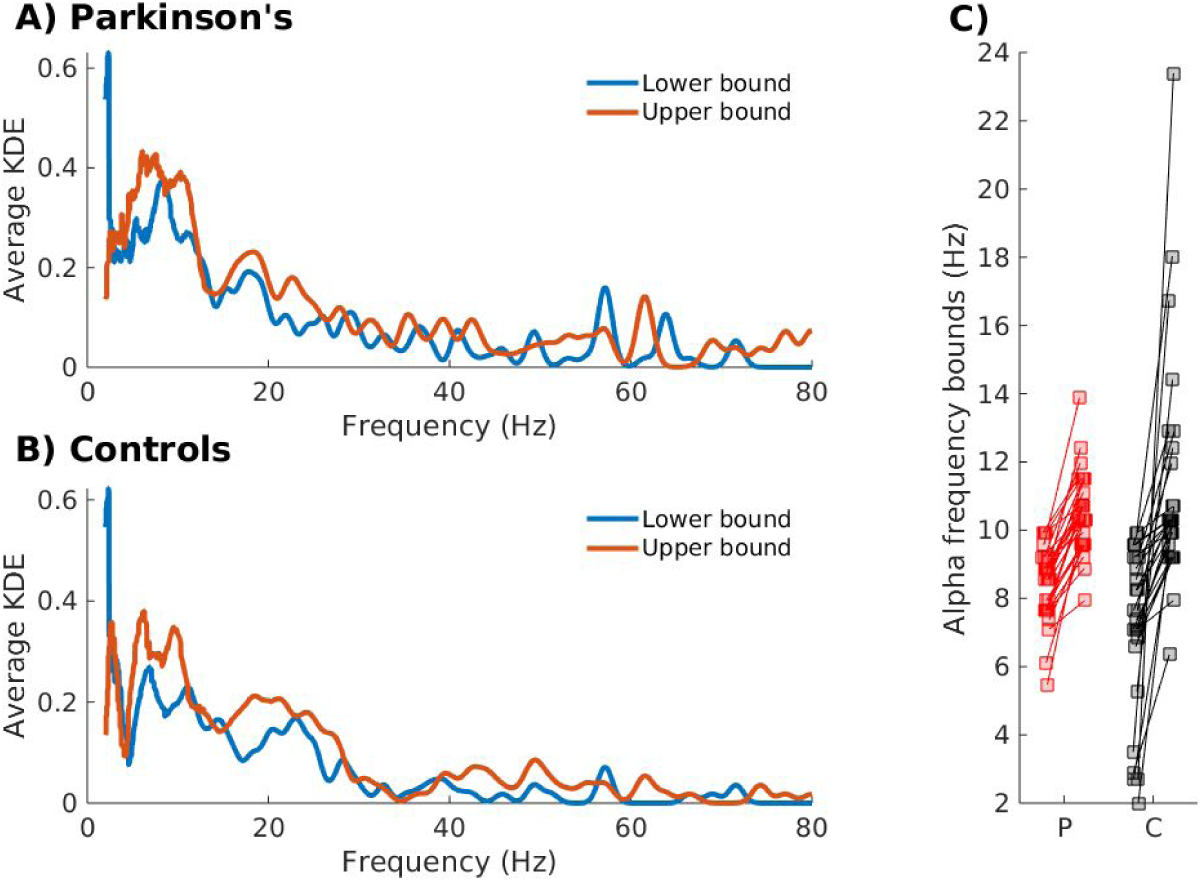
**A-B**) KDEs of frequency boundaries over all participants, separated by group. Higher numbers indicate more consistency in the spectral boundaries across individuals. **C**) Lower and upper alpha band boundaries for the two groups. Each pair of dots is one subject. The outlier control subject is box 899 in Figure S4, and was removed from statistical analysis.

## Discussion

The current and nearly ubiquitous approach of defining frequency bands based on integer boundaries is simple, effective, and reproducible, and it has taken us a long way in linking neural activity to behavior, cognition, and disease. One cannot argue that this approach is invalid or incorrect. However, empirically defined frequency boundaries such as the method presented here opens additional opportunities for explorations and hypothesis-testing of the effects of experiment condition, patient group, genetics, or other factors, on neurophysiological responses.

Existing methods of empirically detecting frequencies rely on the peaks, typically from individual channels (Adam et al., 2014; Chiang et al., 2008; Corcoran et al., 2018; Haegens et al., 2014). These methods have lower sensitivity because they do not leveraging the volume conduction-induced spatial covariance patterns across channels. Furthermore, single-channel approaches suffer from source mixing, particularly in the presence of multiple generators. Nonetheless, the single-channel-peak-detection approach is suitable for identifying the maximum value of a sufficiently prominent peak. The goal here, however, is to identify lower and upper bounds of narrowband activity, even in the absence of robust peak-like deviations from the background spectrum. The increased sensitivity offered by gedBounds allows for detection of frequency ranges that might be closer to the background spectrum.

An advantage of covariance-based methods like gedBounds is that they leverage volume conduction and the spatial autocorrelation structure in the topographies, to increase the signal-to-noise ratio characteristics. Another advantage of gedBounds over single-electrode methods is that it does not rely on the slope of spectral peaks. That is, due to spatial mixing, the downward-slope of the theta peak could overlap with the upward-slope of the alpha peak (e.g., Figure 2a), making them difficult to disentangle at a single electrode. However, as long as the two spectral bands have different anatomical origins (and thus at least slightly different projections onto the electrodes), their corresponding source-separation eigenvectors will decouple the spectral features better than a slope-based or single-channel-based algorithm.

### Multiple components in the same frequency band

The GED for each frequency returns a set of M (channels) eigenvectors that separate the narrowband from the broadband covariance matrices. Although many of these components will capture noise or uninterpretable patterns, it is possible that more than one component per frequency is brain-derived and physiologically meaningful. For example, it is well-known that the brain has multiple generators at the same or similar frequencies in alpha (Olejarczyk et al., 2017; Tognoli and Scott Kelso, 2019) and in theta (Töllner et al., 2017; Zuure et al., n.d.). Focusing exclusively on the component with the largest eigenvalue per frequency is sensible from a statistical perspective, because that top component best separates the narrowband from the broadband activity. But it is also likely that later components are neurally and cognitively meaningful. It would be possible to design an algorithm to pick a component for each frequency, e.g., as maximally responsive to a task condition. One should, however, avoid a selection method based purely on topography, as this can bias the clustering.

### Limitations and considerations

The primary limitation of gedBounds is that it requires parameters to be selected, and different parameters may yield different results. Perhaps the most difficult parameter to select is the epsilon (search step size) for dbscan. A principled approach to selecting this parameter was presented here, but there is no guarantee that that is the optimal approach in all cases. Fortunately, visual inspection of the clusters on the correlation matrix can rule out obviously poor choices for this parameter. Other parameters such as the temporal filter widths may also affect the exact boundaries.

The data used here were taken from resting-state recordings in which there was no cognitive task and therefore no constraints on the time windows to select. If gedBounds is used in a cognitive task, time windows must be selected to create the covariance matrices. This leads to a trade-off between cognitive specificity (using shorter time windows around the cognitive events of interest) and signal quality (using larger time windows to improve the stability of the covariance matrix). The recommendation is to use the widest possible time windows while preserving temporal localization for the cognitive event of interest.

Another limitation is that GED does not distinguish “signal” from “noise”; it simply identifies the spatial pattern that maximally separates two covariance matrices. Therefore, one should not take the gedBounds results *prima facie* without careful inspection of the topographies and time courses. Indeed, the topographies of the component maps in Figure 3 suggests that some of the high- or low-frequency components reflected muscle activity. It is also possible for components to be driven by sensor noise. These are valid results from a statistical perspective, but are not physiologically interpretable.

## Competing or conflicting interests

*none*

## SUPPLEMENTARY MATERIAL

**Figure S1.**
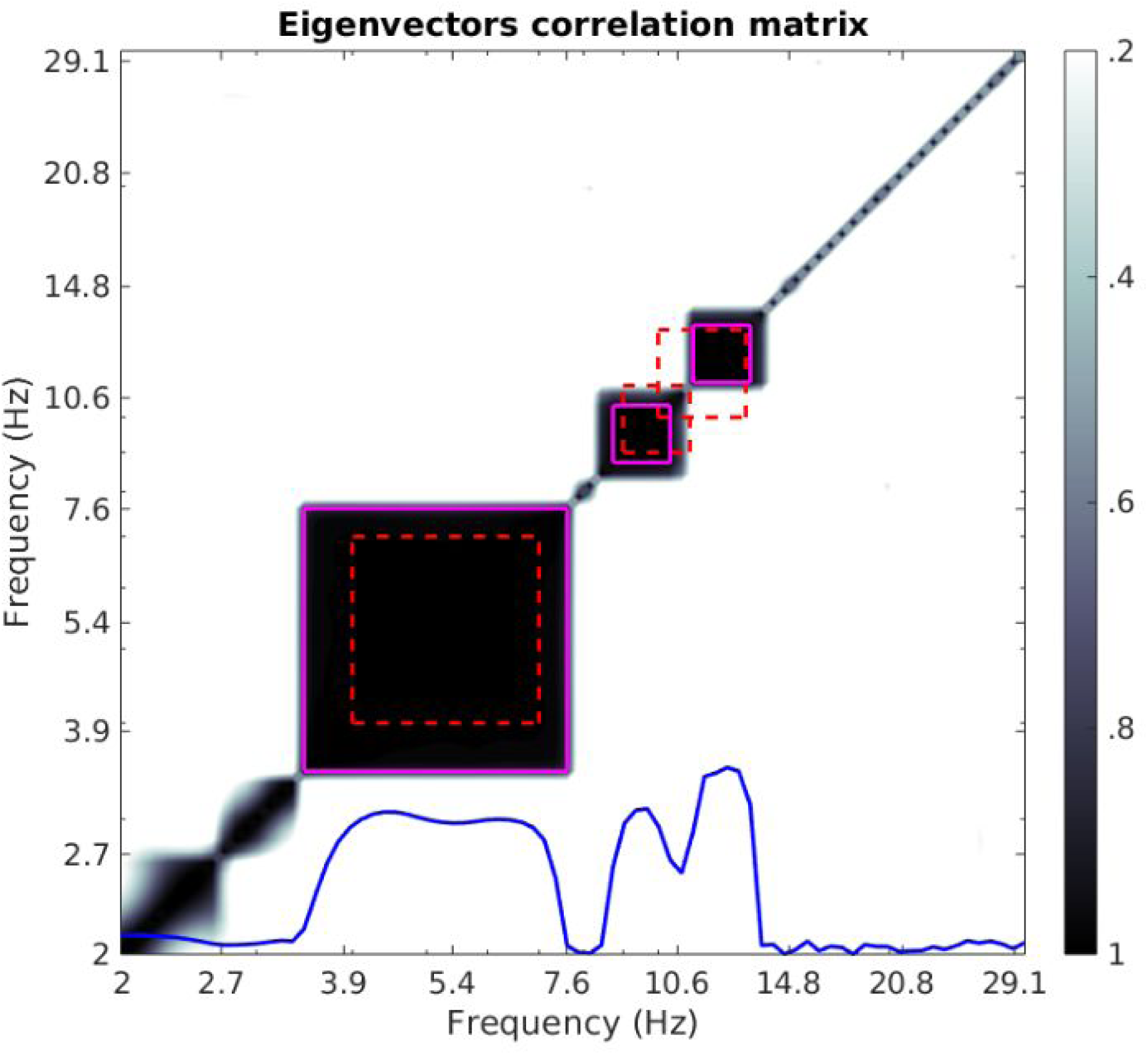
Same as Figure 2a in the main text, but for overlapping frequencies. Note that ground truth frequencies overlapped by a few Hz (red dashed lines), but dbscan identified their non-overlapping spectral regions (solid magenta lines).

**Figure S2.**
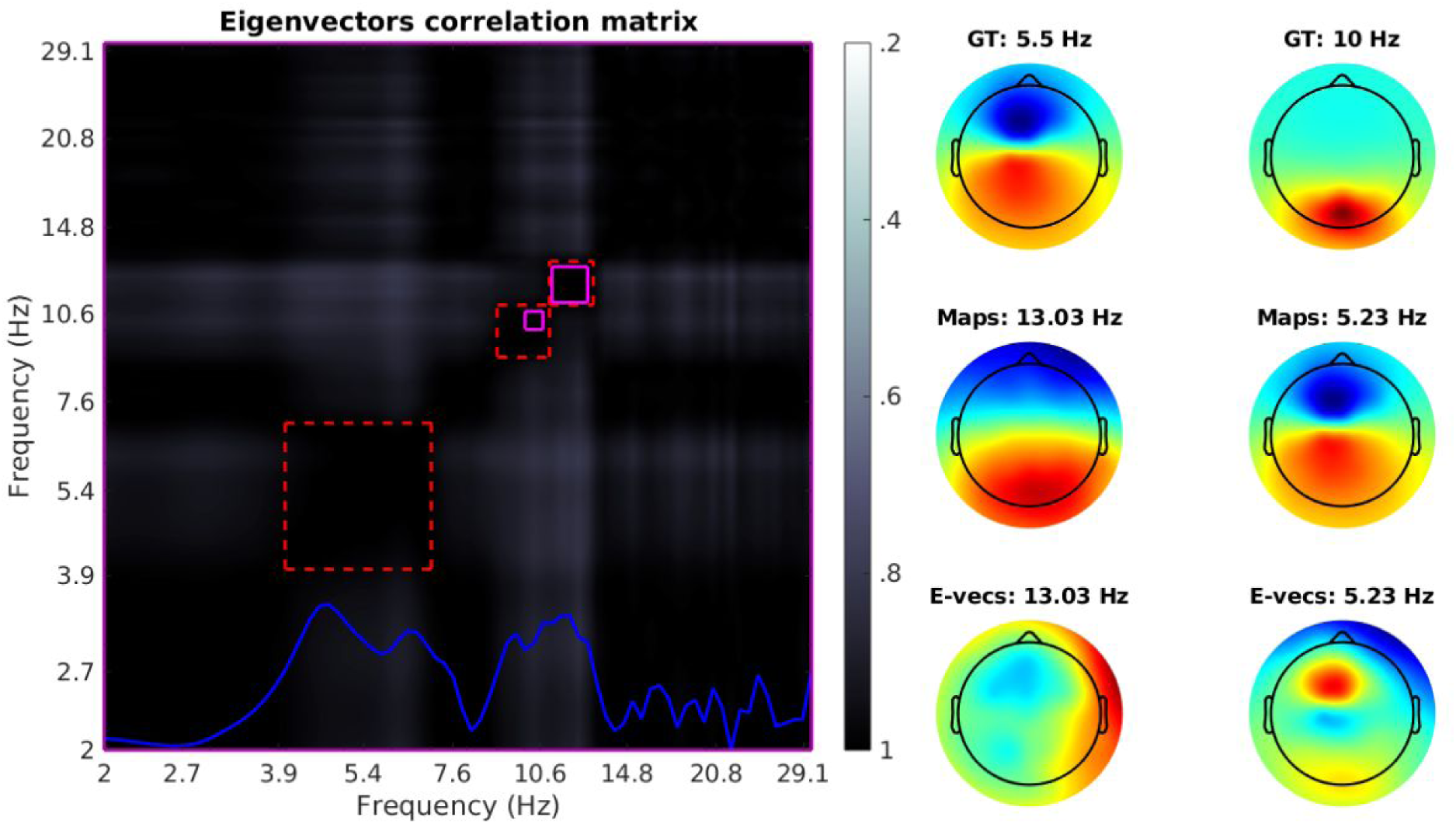
Same as Figure 2 in the main text (including the color scaling), but using PCA on the narrowband covariance matrices instead of GED on narrowband vs. broadband matrices. Data were simulated using the same parameters.

**Figure S3.**
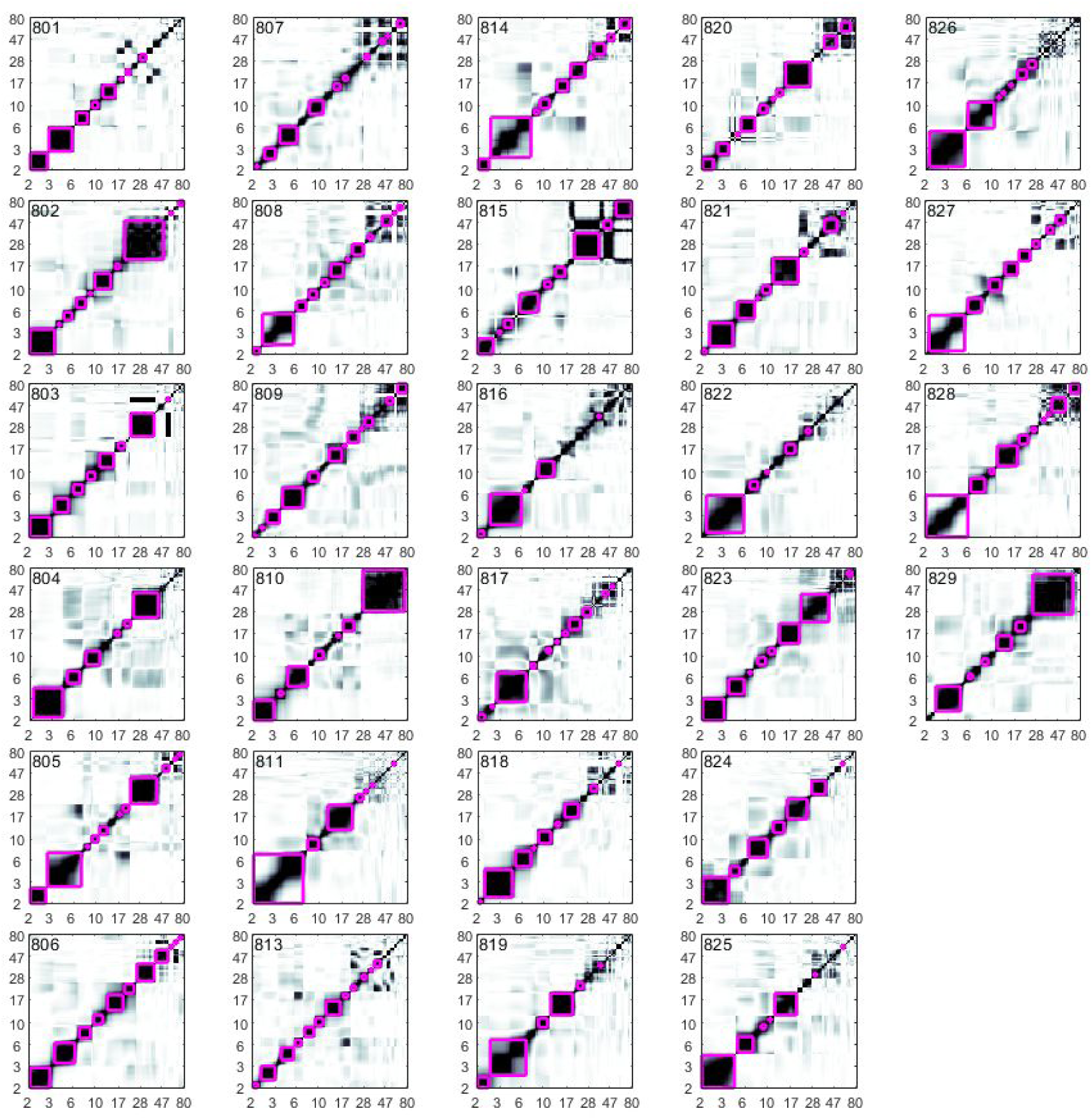
Eigenvector correlation maps with purple boxes indicating dbscan-derived clusters (empirical frequency bands) for all Parkinson’s patients. Numbers outside the boxes indicate frequencies in Hz.

**Figure S4.**
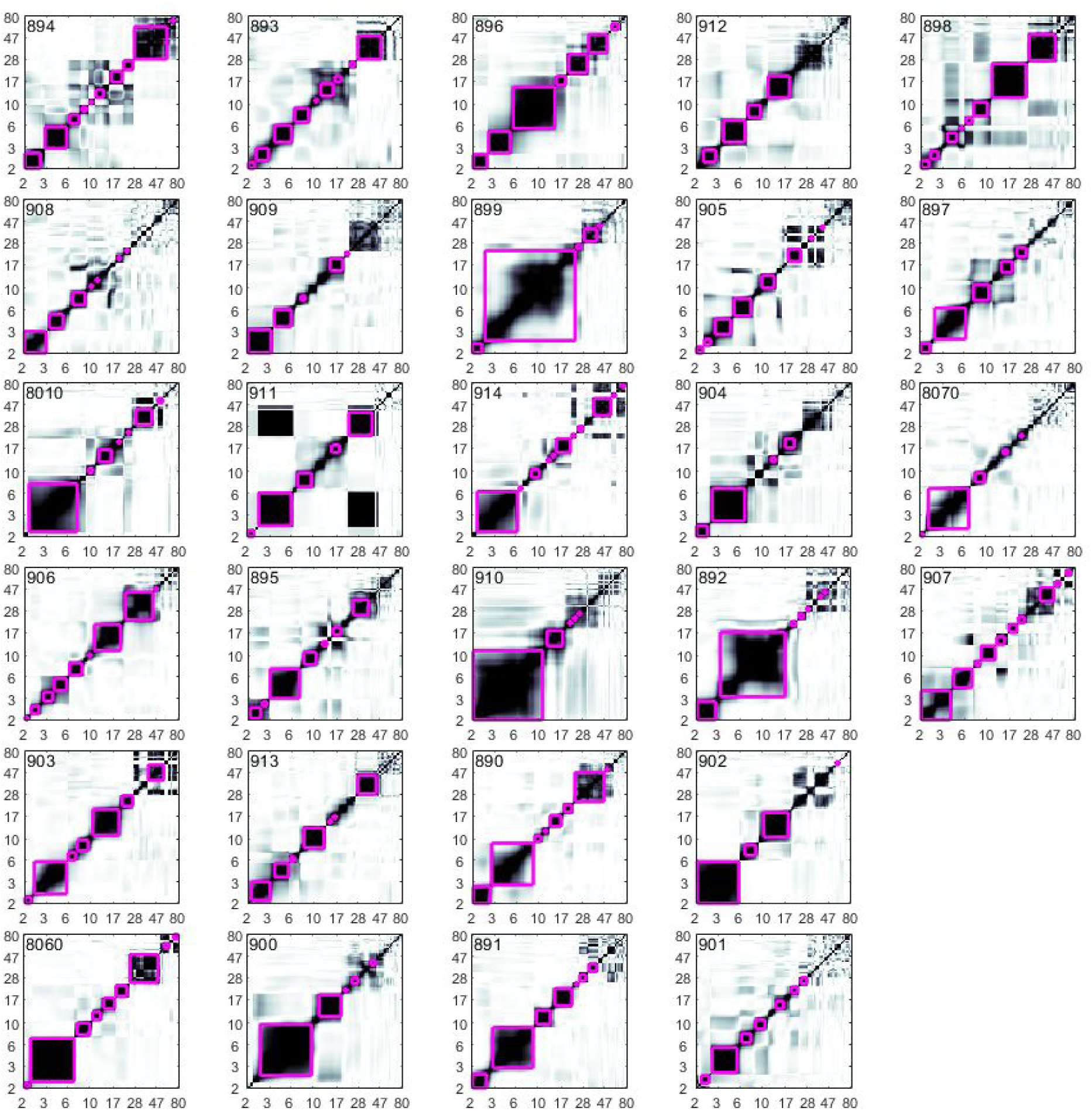
Same as Figure S3 for the control group.

